# AF3Score: A Score-Only Adaptation of AlphaFold3 for Biomolecular Structure Evaluation

**DOI:** 10.1101/2025.05.10.653251

**Authors:** Yu Liu, Qilin Yu, Di Wang, Mingchen Chen

## Abstract

Scoring biomolecular complexes remains central to structural modeling efforts. Recent studies suggest that AlphaFold (AF) – a revolutionary deep learning model for biomolecular structure prediction - has implicitly learned an approximate biophysical energy function. While many researchers highly rely on AF-derived scores for structure evaluation, existing AlphaFold2-based implementations require iterative refinement of the input structure, leading to biased scoring. To address this limitation, we adapted AlphaFold3 into a score-only model, AF3Score, by directly feeding input coordinates into the confidence head while bypassing the diffusion-based structure module. AF3Score demonstrates robust performance in structural quality assessment across diverse systems including monomeric proteins, protein-protein complexes, de novo designed binders, and fold-switching proteins. In benchmarking designed binder screening, AF3Score outperformed state-of-the-art methods for 8 out of 10 targets. Moreover, combining AF3Score with AlphaFold2-derived methods significantly improved the enrichment of experimentally validated binders, increasing the success rate from 15.2% to 31.6%. Additionally, AF3Score effectively identified stable conformations in fold-switching proteins, whereas AlphaFold predominantly predicted only the dominant fold. These findings highlight the broad applicability of AF3Score, from high-throughput screening in de novo binder design to filtering docking-generated poses and molecular dynamics (MD) trajectories.

## Introduction

Recent advances in biomolecular structure prediction have revolutionized drug discovery, enabled rational protein design,^1,2^ and deepened our understanding of molecular interactions. At the forefront of this progress are deep learning-based approaches such as AlphaFold2^3^ and RosettaFold,^4^ which achieved remarkable accuracy by leveraging coevolutionary information from multiple sequence alignments (MSAs). These evolutionary constraints effectively reduce the conformational search space by encoding structural conservation patterns across homologous sequences. However, the predictive performance of these models deteriorates significantly in the absence of MSAs. ^3,5^

The folding landscape of a protein is uniquely determined by its amino acid sequence, with evolutionary pressures sculpting this landscape to ensure proper folding. ^6,7^ While an optimally folded protein requires a funnel-shaped and minimally frustrated energy landscape, functional constraints often introduce local frustrations that hinder efficient conformational search.^8^ This raises an important question: how does AI navigate the rugged folding landscape? One plausible hypothesis is that while AlphaFold2 has learned an accurate energy function for structure evaluation, MSA-derived coevolutionary information is essential for efficiently searching for an approximate global minimum.^9^ This aligns with AlphaFold2’s prediction scheme: an initial MSA-guided search to identify plausible structural templates, followed by iterative refinement using learned energetic potentials.

This concept is further supported by AF2Rank, ^9^ which demonstrated that AlphaFold2’s confidence metrics correlate with structural accuracy even when MSAs are omitted, suggesting the potential of AF2 as a generalizable energy function. Additionally, the metrics show reasonable correlation with experimental success in both monomer and complex structure predictions.^10^

However, existing AlphaFold2-based scoring methods cannot directly evaluate a given structure; instead, they require iterative perturbations of the input coordinates. In contrast, AlphaFold3 exhibits a reduced reliance on MSAs, ^3,5^ while achieving improved prediction accuracy. To harness this learned potential, we developed AF3Score, a score-only evaluator that adapts AlphaFold3 by directly inputting structure coordinates as templates and bypassing the diffusion module. We systematically validated AF3Score across four critical benchmarks: (1) ranking decoys of protein monomers, (2) assessing protein-protein complexes (both antibody and non-antibody), (3) evaluating fold-switching proteins, and (4) assessing binder enrichment in de novo design. Our results demonstrate AF3Score’s superior correlation with experimental outcomes, highlighting its potential as a valuable tool for practical applications and drug discovery.

## Results

### Adapting AlphaFold3 to a score-only model

AF3Score transforms AlphaFold3 into a structure evaluator through two key modifications (Figure 1). First, the genetic search and template search blocks are removed, eliminating reliance on MSAs and external structural databases while preserving the conformer generation component, which is essential for handling ligands and amino acid sidechains. Second, the diffusion module is bypassed, allowing the input coordinates to serve dual roles: they act as structural templates within the template module feeding into the pair representation and as the evaluation target directed to the confidence head. In this process, the atomic coordinates of the input structure are integrated with processed 1D and 2D information from the pairformer to generate quality metrics.

**Figure 1:**
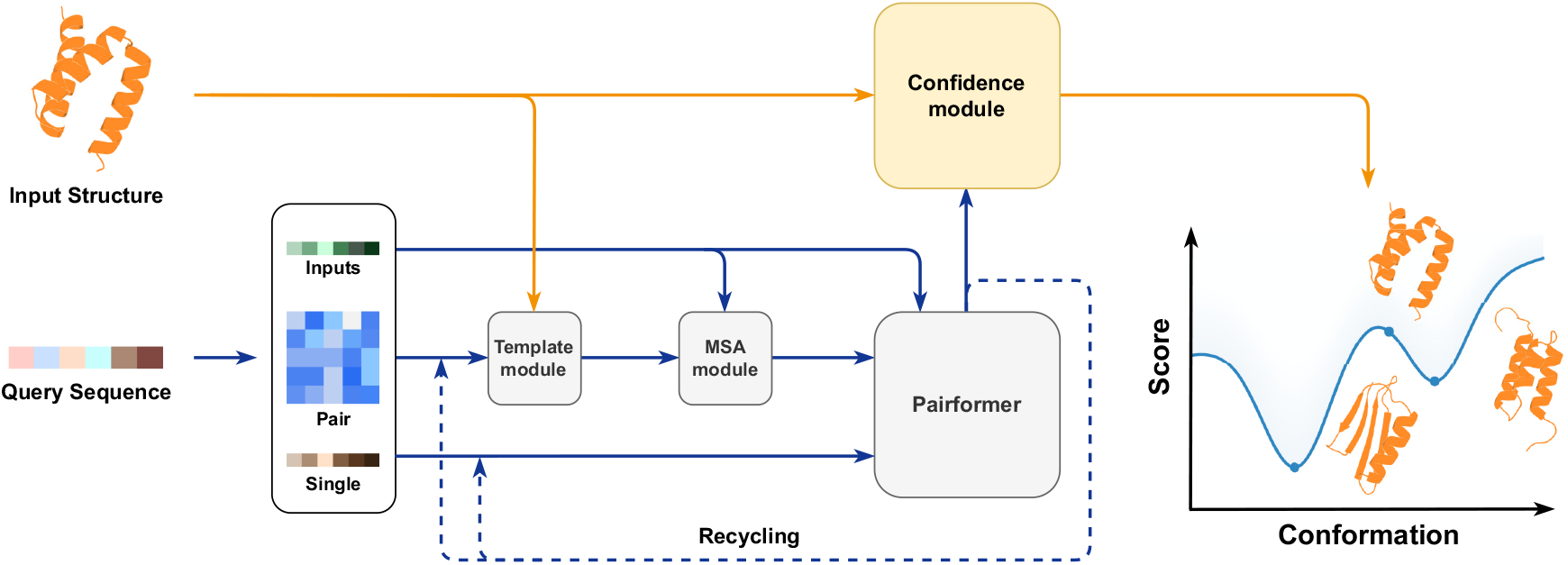
Overview of the AF3Score framework. Schematic representation of the AF3Score pipeline shows processing modules (rectangles) and data flow (arrows) in the streamlined architecture. Yellow arrows indicate the integration of the input structure, while blue arrows represent network activations. The conformational energy landscape was illustrated with the fold-switching protein pairs GA98/GB98 (PDB: 2LHC/2LHD) and a computationally generated decoy from GA98.

This approach differs from AF2Rank, which derives confidence metrics by combining AlphaFold2’s inherent scoring with structural alignment between input and predicted models.

By directly scoring the input structure itself, AF3Score provides a more immediate and unbiased assessment of structural quality.

### Benchmarking AF3Score on Monomers and Protein-Protein Interactions

AlphaFold3 (AF3) demonstrates slightly improved performance in monomeric protein structure prediction compared to AlphaFold2 (AF2). To evaluate whether AF3Score captures a more accurate approximation of the folding landscape, we assessed its ability to rank decoy structures relative to their native conformations. This benchmarking was performed using the Rosetta decoy dataset,^11^ which comprises 133 single-chain targets and approximately 180,000 decoys.

AF3Score was compared against several established methods, including Rosetta, ^12^ DeepAccNet,^13^ and two versions of AF2Rank. ^9^ AF3Score achieved a Spearman correlation of 0.834 with TM-scores across all test cases, surpassing Rosetta (0.760) and DeepAccNet (0.831).

Notably, AF3Score successfully identified near-native structures, with its top-ranked predictions averaging a TM-score of 0.938 (Fig. 2B). These results establish AF3Score as a useful tool for structural quality assessment, although AF2Rank remains slightly superior for this specific task.

**Figure 2:**
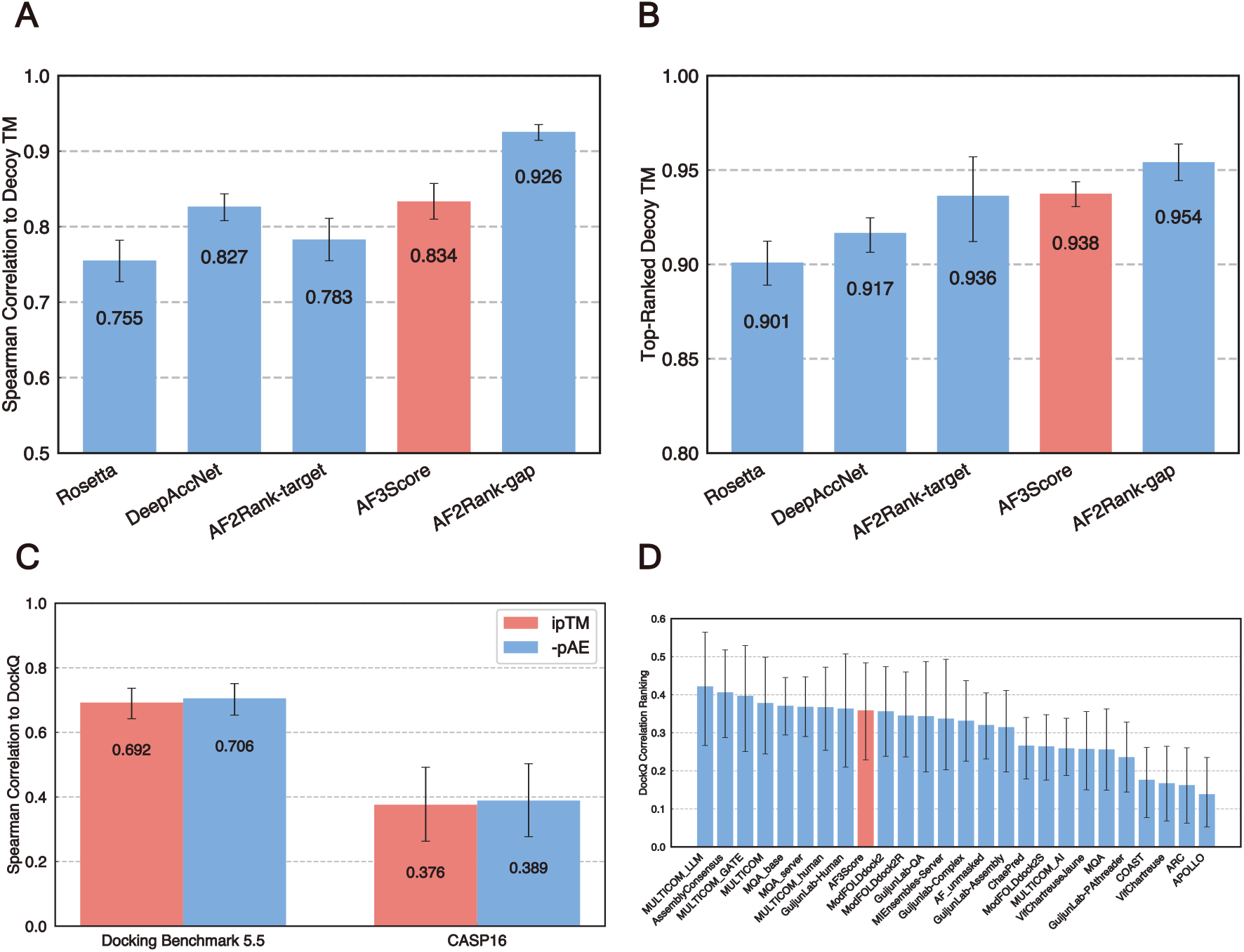
(A,B) Quality assessment of protein structures on the ROSETTA decoy dataset. (A) Spearman correlations between different scoring methods and TM-scores of the decoy docking ensemble. (B) Mean TM-scores of the top-ranked decoys by each method. AF3Score is highlighted as red. (C,D) Evaluations of AF3Score on two PPIs datasets. (C) The spearman correlations between metrics from AF3Score and DockQ values on two datasets are shown. (D) Rankings of assessment of interface quality by 25 methods from CASP16. All error bars represent bootstrap 95% confidence intervals of the mean.

Accurately characterizing protein-protein interactions (PPIs) is essential for understanding cellular functions and drug discovery. Classical computational approaches such as molecular docking^14,15^ play a crucial role in complementing experimental methods, particularly in evaluating interface quality and identifying near-native conformations from computationally generated ensembles.^16,17^ To assess AF3Score’s capability in PPI evaluation, we tested it on two complementary datasets: the Docking Benchmark 5.5,^18^ which covers a diverse range of seen interactions for AlphaFold3, and the CASP16 Quality Assessment (QA) category, which focuses on generalization to novel interactions.^19^ On the Docking Benchmark 5.5 dataset, AF3Score demonstrated strong discrimination of near-native structures against decoy structures from docking simulations (Fig. 2C, Table S2). In CASP16 QA Mode 1 (QMODE1), AF3Score ranked ninth among all participating groups (Fig. 2D, Figures S2, S6, S7), highlighting its robustness in structure-based interaction assessment.

### AF3Score Enhances the Success Rates of De Novo Protein Binder Design

The design of protein binders faces two primary failure modes: (1) the designed sequence fails to adopt its intended monomeric conformation, and (2) the designed sequence correctly folds but does not bind the target.^10^ Previous work showed that AF2-initial-guess offered a promising solution, outperforming physically based methods like Rosetta in predicting experimental successes. ^10^

Using a curated subset from a published dataset of in silico-designed binders, ^10^ AF3Score demonstrated strong discriminative power across ten distinct targets (Fig. 3B,C). Compared to existing methods-including AF2-initial-guess, ^10^ RoseTTAFold2,^20,21^ DeepAccuracyNet,^13^ and Rosetta,^12^ AF3Score achieved superior performance in 6 out of 10 cases. Notably, AF3Score matched or outperformed AF2-initial-guess, the current state-of-the-art for this task, in 8 out of 10 targets (Fig. 3C). A significant improvement was observed when combining AF2-initial-guess with AF3Score, identifying 98 candidates through intersectional selection, with 31 (31.6%) experimentally validated hits. This approach led to a twofold increase in success rate, improving from 15.2% (AF2-initial-guess) to 31.6% (Fig. 3D). AF3Score was further evaluated in a recent competition hosted by Adaptyv Bio, ^22^ which aimed to design binders targeting the epidermal growth factor receptor (EGFR). Compared to AF2, AF3Score demonstrated superior enrichment capabilities for the successful designs. When ranking designs by monomeric metrics, AF3Score identified 75% (3/4) of successful binders within its top 1% ranked designs, whereas AF2 captured only 25% (1/4) (Fig. 3E). Using complex metrics, the iPTM from AF3Score achieved a validation accuracy of 50% (2/4), whereas AF2’s PAE interaction metric failed to identify any true binders (0/4) (Fig. 3F). Notably, a consensus filtering approach that combined the top 5% of candidates from both AF3Score and AF2 achieved a success rate of 55.6% (5/9 validated).

**Figure 3:**
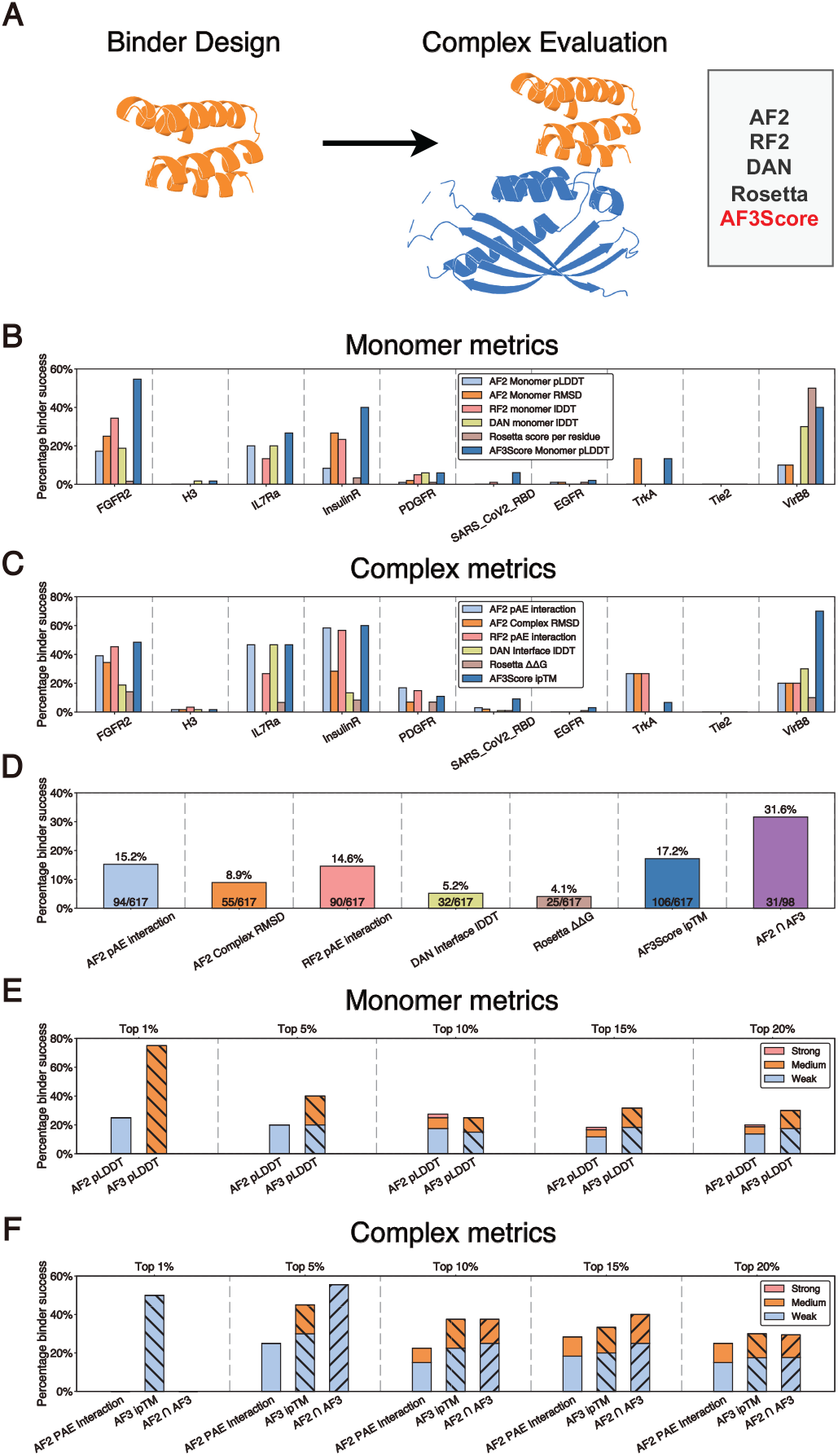
Benchmarking on the EGFR binder design contest by Adaptyv Bio. (A) Schematic illustration of different methods used to assess designed protein complexes, with mini-binders shown in yellow and target proteins in blue. (B, C, D) Evaluation of designed protein binders across 10 targets from Cao et al. (B, C) Comparison of experimental success rates (YSD SC50 *<* 4 *µ*M) for the top 1% of designs selected based on different monomeric (B) or protein complex (C) metrics. (D) Overall success rates of the top 1% of designs selected using individual protein complex metrics, as well as the intersection of AF2 pAE interaction and AF3Score ipTM (AF2 ∩ AF3). (E, F) Evaluation of designed protein binders targeting EGFR. Experimental validation identified a success rate of 13.25% (53/400 designs). (E, F) Comparison of experimental success rates for designs selected based on different monomeric (E) or protein complex (F) metrics. Bars indicate the percentage of successful binders within different binding strength categories (strong, medium, and weak) across the top 1%, 5%, 10%, 15%, and 20% of structures ranked by each metric.

### Benchmarking AF3Score on Fold-Switching Proteins

Fold-switching proteins adopt two or more distinct folded states from a single amino acid sequence, undergoing either spontaneous conformational interconversion or stimulus-responsive structural transitions.^23,24^ Predicting alternative folds remains a major challenge for AlphaFold2 (AF2).^25^ A recent benchmark^26^ evaluating all AlphaFold versions, including AF2,^3^ AF2 multimer,^27^ AF3,^5^ and enhanced sampling methods,^28,29^ demonstrated AF2’s limitations in predicting fold-switching events. While multiple sequence alignments (MSAs) provide valuable evolutionary context for dominant protein structures, they often fail to capture alternative conformations. ^28,30^ For a comprehensive scoring function, both dominant and alternative conformations should achieve high confidence scores. To test this hypothesis, we reframed fold-switching prediction as a structural scoring problem.

Using an established database of fold-switching proteins,^25^ we curated 45 protein pairs, excluding cases with insertions, deletions, or mutations in their fold-switching regions. Initial evaluation using AF3Score (Fig. 4A) revealed comparable confidence scores for both conformations, with mean values of 83.15 and 81.85 for Fold 1 and Fold 2, respectively, unlike existing methods, which often exhibit strong bias toward a single conformation.

**Figure 4:**
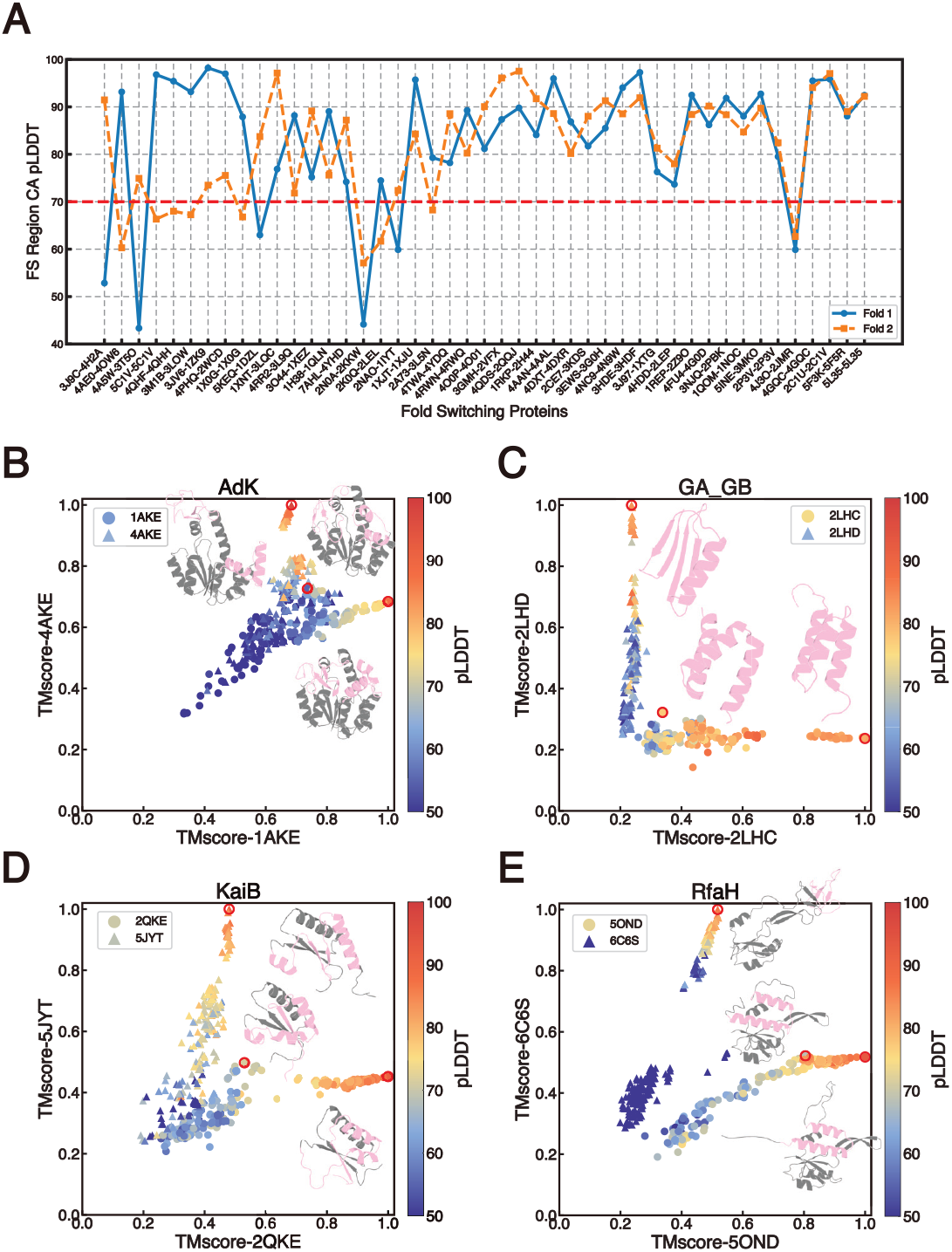
Benchmarking AF3Score on fold-switching proteins. (A) Confidence scores assigned by AF3Score for 45 pairs of fold-switching proteins. The two native conformations are colored in orange and blue. The red line indicates a fold-switching (FS) region pLDDT threshold of 70, marking high-confidence predictions in these regions. (B-E) Analysis of the representative fold-switching proteins. The x- and y-axes represent TM-scores relative to each native fold. Each point corresponds to a structure, colored according to the C*α* pLDDT score of its fold-switching region. Triangles and circles distinguish between the two alternative folds. Native structures and selected decoys (highlighted by red circles) are shown, with fold-switching regions in pink and the remaining structure in grey.

To further assess AF3Score’s ability to capture fold-switching, we examined the conformational landscapes of several well-characterized fold-switching proteins (Fig. 4B-E and Figure S12). While auxiliary approaches, such as local energetic frustration analysis, can help AF2 sample alternative states,^31^ we investigated whether AF3Score inherently overcomes this limitation. For each of the six cases, 200 structures were sampled using 3D-Robot between the two alternate states, and the resulting structure ensemble was later scored.

A particularly illustrative example is Adenylate kinase (AdK) (Fig. 4B), a multidomain protein that serves as a model for allosteric regulation.^31^ While certain decoy structures exhibited moderate structural similarity to both native folds (TM-score *>* 0.7), their AF3Score confidence values were notably low, highlighting its sensitivity to fold-specific features. Similar trend was also observed for KaiB (Fig. 4D), RfaH (Fig. 4E) and other proteins (Fig. S12).

## Conclusion

Building upon the biophysical insights captured in AlphaFold3’s energy function, we introduce AF3Score, a scoring framework for evaluating biomolecular structure. This scoreonly model demonstrates broad applicability across multiple tasks, including the assessment of monomeric protein, protein-protein interaction, fold-switching protein, and de novo designed protein binders.

Our results show that AF3Score enhances success rates in de novo protein design compared to state-of-the-art methods. By assessing the input sequence against a provided structure rather than a predicted one, AF3Score provides a more robust estimation of structural compatibility. This makes AF3Score a powerful tool for downstream screening in binder design, efficiently prioritizing candidates by assessing both monomer stability and target binding.

## Data Availability Statement

The code is available on GitHub https://github.com/Mingchenchen/AF3Score/ and is installable. Full documentation complete with worked examples of how the package can be used are also available at github.

## References

(1) Watson, J. L. et al. De Novo Design of Protein Structure and Function with RFdiffusion. Nature 2023, 620, 1100–.

(2) Pacesa, M. et al. BindCraft: One-Shot Design of Functional Protein Binders. 2024.

(3) Jumper, J. et al. Highly Accurate Protein Structure Prediction with AlphaFold. Nature 2021, 596, 589–.

(4) Baek, M. et al. Accurate Prediction of Protein Structures and Interactions Using a Three-Track Neural Network. Science 2021, 373, 876–.

(5) Abramson, J. et al. Accurate Structure Prediction of Biomolecular Interactions with AlphaFold 3. Nature 2024, 630, 500–.

(6) Schafer, N. P.; Kim, B. L.; Zheng, W.; Wolynes, P. G. Learning To Fold Proteins Using Energy Landscape Theory. Israel Journal of Chemistry 2014, 54, 1337–.

(7) Wolynes, P. G. Evolution, Energy Landscapes and the Paradoxes of Protein Folding. Biochimie 2015, 119, 230–.

(8) Chen, M.; Chen, X.; Schafer, N. P.; Clementi, C.; Komives, E. A.; Ferreiro, D. U.; Wolynes, P. G. Surveying Biomolecular Frustration at Atomic Resolution. Nature Communications 2020, 11, 5944.

(9) Roney, J. P.; Ovchinnikov, S. State-of-the-Art Estimation of Protein Model Accuracy Using AlphaFold. Physical Review Letters 2022, 129, 238101.

(10) Bennett, N. R.; Coventry, B.; Goreshnik, I.; Huang, B.; Allen, A.; Vafeados, D.; Peng, Y. P.; Dauparas, J.; Baek, M.; Stewart, L.; DiMaio, F.; De Munck, S.; Savvides, S. N.; Baker, D. Improving de Novo Protein Binder Design with Deep Learning. Nature Communications 2023, 14, 2625.

(11) Park, H.; Bradley, P.; Greisen, P.; Liu, Y.; Mulligan, V. K.; Kim, D. E.; Baker, D.; DiMaio, F. Simultaneous Optimization of Biomolecular Energy Functions on Features from Small Molecules and Macromolecules. Journal of Chemical Theory and Computation 2016, 12, 6212–.

(12) Alford, R. F. et al. The Rosetta All-Atom Energy Function for Macromolecular Modeling and Design. Journal of Chemical Theory and Computation 2017, 13, 3048–.

(13) Hiranuma, N.; Park, H.; Baek, M.; Anishchenko, I.; Dauparas, J.; Baker, D. Improved Protein Structure Refinement Guided by Deep Learning Based Accuracy Estimation. Nature Communications 2021, 12, 1340.

(14) Yan, Y.; Tao, H.; He, J.; Huang, S.-Y. The HDOCK Server for Integrated Protein– Protein Docking. Nature Protocols 2020, 15, 1852–.

(15) Honorato, R. V.; Trellet, M. E.; Jimenez-Garcia, B.; Schaarschmidt, J. J.; Giulini, M.; Reys, V.; Koukos, P. I.; Rodrigues, J. P. G. L. M.; Karaca, E.; Van Zundert, G. C. P.; Roel-Touris, J.; Van Noort, C. W.; Jandova, Z.; Melquiond, A. S. J.; Bonvin, A. M. J. J. The HADDOCK2.4 Web Server for Integrative Modeling of Biomolecular Complexes. Nature Protocols 2024, 19, 3241–.

(16) Vajda, S.; Hall, D. R.; Kozakov, D. Sampling and Scoring: A Marriage Made in Heaven: Sampling and Scoring. Proteins: Structure, Function, and Bioinformatics 2013, 81, 1884–.

(17) Cao, Y.; Shen, Y. Energy-based Graph Convolutional Networks for Scoring Protein Docking Models. Proteins: Structure, Function, and Bioinformatics 2020, 88, 1099–.

(18) Vreven, T.; Moal, I. H.; Vangone, A.; Pierce, B. G.; Kastritis, P. L.; Torchala, M.; Chaleil, R.; Jimenez-Garcia, B.; Bates, P. A.; Fernandez-Recio, J.; Bonvin, A. M.; Weng, Z. Updates to the Integrated Protein–Protein Interaction Benchmarks: Docking Benchmark Version 5 and Affinity Benchmark Version 2. Journal of Molecular Biology 2015, 427, 3041–.

(19) Home - CASP16. https://predictioncenter.org/casp16/index.cgi.

(20) Baek, M.; McHugh, R.; Anishchenko, I.; Jiang, H.; Baker, D.; DiMaio, F. Accurate Prediction of Protein–Nucleic Acid Complexes Using RoseTTAFoldNA. Nature Methods 2024, 21, 121–.

(21) Watson, J. L. et al. Broadly Applicable and Accurate Protein Design by Integrating Structure Prediction Networks and Diffusion Generative Models. 2022.

(22) Adaptyv Foundry. https://foundry.adaptyvbio.com/competition.

(23) Porter, L. L.; Looger, L. L. Extant Fold-Switching Proteins Are Widespread. Proceedings of the National Academy of Sciences 2018, 115, 5973–.

(24) Schafer, J. W.; Porter, L. L. Evolutionary Selection of Proteins with Two Folds. Nature Communications 2023, 14, 5478.

(25) Porter, L. N. E. AlphaFold2 Fails to Predict Protein Fold Switching.

(26) Chakravarty, D.; Schafer, J. W.; Chen, E. A.; Thole, J. F.; Ronish, L. A.; Lee, M.; Porter, L. L. AlphaFold Predictions of Fold-Switched Conformations Are Driven by Structure Memorization. Nature Communications 2024, 15, 7296.

(27) Evans, R. et al. Protein Complex Prediction with AlphaFold-Multimer. 2021.

(28) Stein, R. A.; Mchaourab, H. S. SPEACH AF: Sampling Protein Ensembles and Conformational Heterogeneity with Alphafold2. PLOS Computational Biology 2022, 18, e1010483.

(29) Wayment-Steele, H. K.; Ojoawo, A.; Otten, R.; Apitz, J. M.; Pitsawong, W.; Homberger, M.; Ovchinnikov, S.; Colwell, L.; Kern, D. Predicting Multiple Conformations via Sequence Clustering and AlphaFold2. Nature 2024, 625, 839–.

(30) Kalakoti, Y.; Wallner, B. AFsample2: Predicting Multiple Conformations and Ensembles with AlphaFold2. 2024.

(31) Guan, X.; Tang, Q.-Y.; Ren, W.; Chen, M.; Wang, W.; Wolynes, P. G.; Li, W. Predicting Protein Conformational Motions Using Energetic Frustration Analysis and AlphaFold2. Proceedings of the National Academy of Sciences 2024, 121, e2410662121.

