## Supporting Information for "AF3Score: A Score-Only Adaptation of AlphaFold3 for Biomolecular Structure Evaluation"

#### Limitations of AF2-initial-guess and advantages of AF3Score in structural evaluation

One critical limitation of the AF2-initial-guess, is the discrepancy between evaluating the input and output structures. AF2-initial-guess initializes the AF2 pair representation with the designed complex structure but then generates a predicted structure along with its corresponding metrics. Because these metrics are based on the predicted structure rather than the original input, they do not fully reflect the designed complex. In contrast, AF3Score evaluates the input sequence against the provided structure, potentially offering a more robust estimation of input conformations.

#### AF3Score accurately identifies diverse binding modes and epitopes in case studies

In the complexes formed between TEM-1 $\beta$-lactamase and $\beta$-lactamase inhibitor protein (BLIP) or $\beta$-lactamase inhibitor protein-II (BLIP-II), although BLIP and BLIP-II adopt completely different folds, they both exhibit a competitive mode of inhibition toward TEM-1.[1] In this case, AF3Score accurately recognized the binding modes despite the differences in their binding modes (Supplementary Figures S8A and S8B). Another example illustrates the ability of AF3Score to identidy distinct epitopes of the same antigen. Interleukin-1$\beta$ (IL-1$\beta$) is a key orchestrator in inflammatory and immune responses.[2] Two highly specific IL-1$\beta$ monoclonal antibodies, Canakinumab and gevokizumab, exhibit divergent binding mechanisms. Canakinumab competitively inhibits IL-1$\beta$ association with its receptor due to steric overlap between their binding interfaces on IL-1$\beta$. In contrast, gevokizumab occupies an allosteric site on IL-1$\beta$, forming a complex that reduces the binding affinity of IL-1RI. In this case, AF3Score accurately identified both epitopes, demonstrating the capability of AF3Score to recognize different local structural information and thereby distinguish between distinct binding modes (Supplementary Figures S8C).

Despite its reasonable performance, AF3Score faces challenges in capturing certain novel interfaces, and this might be due to a lack of more interaction data during training. For example, in the caspase-9:BIR3 complex, the confidence scores of docking decoys fall within a narrow range (less than 0.2), and the native structure's score is only marginally higher than those of the decoys (Supplementary Fig. S9).

#### The moderate performance of Af3Score on protein-ligand complexes

We further extended AF3Score to evaluate protein-ligand complexes, where it exhibited moderate discriminative power (Supplementary Fig. S15A). A limitation is its reduced accuracy in ranking near-native ligand poses, likely due to the lack of template-guided refinement for ligand coordinates during the representation updates in AlphaFold3 (Supplementary Fig. S15B). Our preliminary control experiments, where templates of proteins were intentionally excluded, demonstrated that the ability of AF3Score to assess structural plausibility was significantly impaired.

### Methods

#### Dataset of protein-protein interactions

We selected 188 protein complexes from Docking Benchmark 5.5 that do not contain cofactors or ligands,[3] of which 63 structures are antigen-antibody complexes. We used HDOCK to perform protein-protein docking simulations on the bound structures of these complexes. The docking process utilized HDOCK's default parameter settings, generating 100 candidate conformations for each complex. The quality of each docked pose was evaluated using DockQ v2,[4] focusing specifically on the interface DockQ scores between receptor and ligand chains.

From the 38 targets in CASP16's QMODE1,[5] we selected 30 targets for testing, including 13 homo-oligomers and 17 hetero-oligomers. The selection criteria were: (1) targets with fewer than 3072 amino acids to avoid memory limitations, (2) targets with determined stoichiometry, and (3) targets with available EMA analysis results. We evaluated all models for the 30 targets, although only models with index 1 were considered for QMODE1 ranking.

#### Dataset of de novo designed binders

From a published dataset of approximately 700,000 minibinders,[6] we extracted all 1,303 experimentally validated successful binders, and then randomly sampled 61,781 non-binding designs to constitute our dataset. For the ten minibinder structures with available crystal structures, we used HDOCK to generate 100 decoy structures for each, and evaluated their interface quality using DockQ v2 scores between the designed binder and target protein chains.

#### Dataset of fold switching proteins

We curated 45 fold-switching proteins from a dataset of 98 fold-switching proteins.[7] Among the original 98 proteins, 93 had their alternate conformations solved in PDB and were considered in this study. For each fold-switching protein, we examined its dominant and alternative conformations, excluding those with insertions, deletions, or mutations in the fold-switching region to ensure identical sequences in this region between the two conformations, resulting in 45 proteins. The decoy generation was performed separately using both the dominant and alternative conformations as references. For each target conformation, we generated 200 decoy structures using 3DRobot with an RMSD cutoff of 12Å. 3DRobot first identifies structure scaffolds from a non-redundant PDB library, then performs structure assembly simulations based on each scaffold, and finally selects decoys through a two-step energy minimization procedure.

#### Dataset of protein-ligand interactions

We selected 178 protein-ligand complexes from the posebusters dataset for our study.[8] To ensure data quality, we implemented a comprehensive preprocessing pipeline. First, we removed common crystallants (including SO4, GOL, EDO, PO4, ACT, PEG, etc.) from the original PDB files, as these molecules are crystallization artifacts rather than natural binding partners. Subsequently, we standardized the structures by first converting ligands from SDF to PDB format, then standardizing ligand atom nomenclature according to the Chemical Component Dictionary, and reassigning residue numbers and chain IDs. To generate decoy poses, we employed AutoDock Vina with an exhaustiveness of 32 and a search box of 15Å centered on the crystallographic ligand position, generating 50 docked poses for each complex. The LRMSD value of each docked pose was calculated relative to the pose of the native binding pose by DockQ v2.[4]

### References

(1) Lim, D.; Park, H. U.; De Castro, L.; Kang, S. G.; Lee, H. S.; Jensen, S.; Lee, K. J.; Strynadka, N. C. J. Crystal Structure and Kinetic Analysis of $\beta$-Lactamase Inhibitor

Protein-II in Complex with TEM-1 $\beta$-Lactamase. *Nature Structural Biology* **2001**, *8*, 848–852.

(2) One Target—Two Different Binding Modes: Structural Insights into Gevokizumab and Canakinumab Interactions to Interleukin-$\beta$. *Journal of Molecular Biology* **2013**, *425*, 94–111.

(3) Vreven, T.; Moal, I. H.; Vangone, A.; Pierce, B. G.; Kastritis, P. L.; Torchala, M.; Chaleil, R.; Jiménez-García, B.; Bates, P. A.; Fernandez-Recio, J.; Bonvin, A. M.; Weng, Z. Updates to the Integrated Protein–Protein Interaction Benchmarks: Docking Benchmark Version 5 and Affinity Benchmark Version 2. *Journal of Molecular Biology* **2015**, *427*, 3031–3041.

(4) Mirabello, C.; Wallner, B. DockQ v2: Improved Automatic Quality Measure for Protein Multimers, Nucleic Acids, and Small Molecules. *Bioinformatics* **2024**, *40*, btae586.

(5) Home - CASP16. https://predictioncenter.org/casp16/index.cgi.

(6) Cao, L. et al. Design of Protein-Binding Proteins from the Target Structure Alone. *Nature* **2022**, *605*, 551–560.

(7) Porter, L. N. E. AlphaFold2 Fails to Predict Protein Fold Switching.

(8) Buttenschoen, M.; Morris, G. M.; Deane, C. M. PoseBusters: AI-based Docking Methods Fail to Generate Physically Valid Poses or Generalise to Novel Sequences. *Chemical Science* **2024**, *15*, 3130–3139.

**Algorithm 1:** AF3Score workflow

**Input:** A protein monomer or complex structure in PDB format
**Output:** Quality assessment scores (ipTM, PAE, pLDDT, pTM)
**1. Structure Processing:**;
Extract and save individual chains as CIF files;
Generate JSON configuration using CIF files as templates;
**2. Coordinate Conversion:**;
Convert PDB file to JAX arrays with AlphaFold's atom14 representation;
Pad to nearest bucket size (256-3072) and save as H5 format;
**3. AF3 Inference:**;
Run AF3 inference with atomic coordinates in H5 format;
Extract quality assessment scores;
**return** *quality assessment scores*

Table 1: Spearman Correlation and Top-ranked Metrics of AF3Score Metrics for the Rosetta dataset. After testing multiple scoring options (pTM, pAE, pLDDT, and their combinatorial forms), we chose pTM as the confidence score, since it best matched TM-scores, which aligns with the ranking score used in AF3.

| AF3Score Metrics | Correlation | Top-ranked |
|---|---|---|
| pLDDT | $0.808 \pm 0.023$ | $0.920 \pm 0.020$ |
| pTM | $0.834 \pm 0.024$ | $\mathbf{0.938 \pm 0.007}$ |
| PAE | $0.833 \pm 0.024$ | $0.934 \pm 0.011$ |
| pLDDT x pTM | $\mathbf{0.835 \pm 0.020}$ | $0.938 \pm 0.012$ |

Table 2: Pearson correlations between AF3Score metrics and DockQ scores on the Docking Benchmark 5.5 dataset, comprising 188 protein complexes (63 antigen-antibody and 125 other protein-protein interactions).

| Metric | All (N=188) | AbAg (N=63) | Other (N=125) |
|---|---|---|---|
| ipTM | $0.692 \pm 0.047$ | $0.717 \pm 0.074$ | $0.680 \pm 0.058$ |
| pTM | $0.672 \pm 0.049$ | $0.687 \pm 0.072$ | $0.664 \pm 0.064$ |
| pLDDT | $0.470 \pm 0.060$ | $0.496 \pm 0.092$ | $0.457 \pm 0.073$ |
| PAE | $\mathbf{0.706 \pm 0.047}$ | $\mathbf{0.731 \pm 0.074}$ | $\mathbf{0.693 \pm 0.062}$ |

Table 3: Spearman correlations between AF3Score metrics and TMscore or DockQ for multimer targets in QA. In CASP16 Quality Assessment Mode 1 (QMODE1), only the model labeled "1" (model index 1) was used. "All Models" indicates that the analysis included models with all indices.

| | QMODE1 | | All models | |
|---|---|---|---|---|
| | TMscore | DockQ | TMscore | DockQ |
| pLDDT | $0.349 \pm 0.113$ | $0.34 \pm 0.116$ | $0.359 \pm 0.101$ | $0.374 \pm 0.110$ |
| pTM | $0.365 \pm 0.103$ | $0.341 \pm 0.122$ | $0.372 \pm 0.091$ | $0.369 \pm 0.109$ |
| ipTM | $0.356 \pm 0.103$ | $0.35 \pm 0.127$ | $0.364 \pm 0.098$ | $0.376 \pm 0.107$ |
| PAE | $\mathbf{0.393 \pm 0.103}$ | $\mathbf{0.359 \pm 0.119}$ | $\mathbf{0.390 \pm 0.096}$ | $\mathbf{0.389 \pm 0.106}$ |

Table 4: Mean TMscore and DockQ for top-ranked models for each target selected by different metrics of AF3Score. AF3Score maintained its discriminative power in selecting high-quality models. The analysis included models with all indices for each multimer target in QA.

| | Homomultimers | | Heteromultimers | |
|---|---|---|---|---|
| | TMscore | DockQ | TMscore | DockQ |
| pLDDT | $0.775 \pm 0.114$ | $0.519 \pm 0.183$ | $0.835 \pm 0.078$ | $\mathbf{0.584 \pm 0.116}$ |
| pTM | $0.877 \pm 0.096$ | $0.581 \pm 0.151$ | $0.841 \pm 0.078$ | $0.576 \pm 0.110$ |
| ipTM | $\mathbf{0.905 \pm 0.073}$ | $\mathbf{0.585 \pm 0.144}$ | $0.841 \pm 0.080$ | $0.575 \pm 0.118$ |
| PAE | $0.802 \pm 0.110$ | $0.523 \pm 0.172$ | $\mathbf{0.844 \pm 0.066}$ | $0.565 \pm 0.094$ |

Table 5: Performance comparison of monomer and complex-based metrics from AF3Score in selecting successful designs across 10 protein targets.

| Target | Monomer | | | Complex | | | | |
|---|---|---|---|---|---|---|---|---|
| | pLDDT | PAE | pTM | pLDDT | PAE_interaction | PAE_interface | pTM | ipTM |
| FGFR2 | 54.69% | 43.75% | 48.44% | 46.88% | 51.56% | 45.31% | 51.56% | 48.44% |
| H3 | 1.69% | 0.00% | 0.00% | 1.69% | 1.69% | 1.69% | 1.69% | 1.69% |
| IL7Ra | 26.67% | 33.33% | 26.67% | 20.00% | 40.00% | 46.67% | 26.67% | 46.67% |
| InsulinR | 40.00% | 31.67% | 38.33% | 33.33% | 50.00% | 40.00% | 73.33% | 60.00% |
| PDGFR | 5.94% | 5.94% | 9.90% | 4.95% | 11.88% | 12.87% | 7.92% | 10.89% |
| SARS_CoV2_RBD | 6.06% | 1.01% | 3.03% | 6.06% | 9.09% | 9.09% | 9.09% | 9.09% |
| EGFR | 2.04% | 0.00% | 3.06% | 1.02% | 1.02% | 1.02% | 1.02% | 3.06% |
| TrkA | 13.33% | 6.67% | 26.67% | 13.33% | 13.33% | 13.33% | 6.67% | 6.67% |
| Tie2 | 0.00% | 0.00% | 0.00% | 0.00% | 0.00% | 0.00% | 0.00% | 0.00% |
| VirB8 | 40.00% | 20.00% | 30.00% | 50.00% | 70.00% | 50.00% | 70.00% | 70.00% |
| Overall | **19.04%** | 14.24% | 18.61% | 17.73% | 24.86% | 22.00% | 24.80% | **25.65%** |

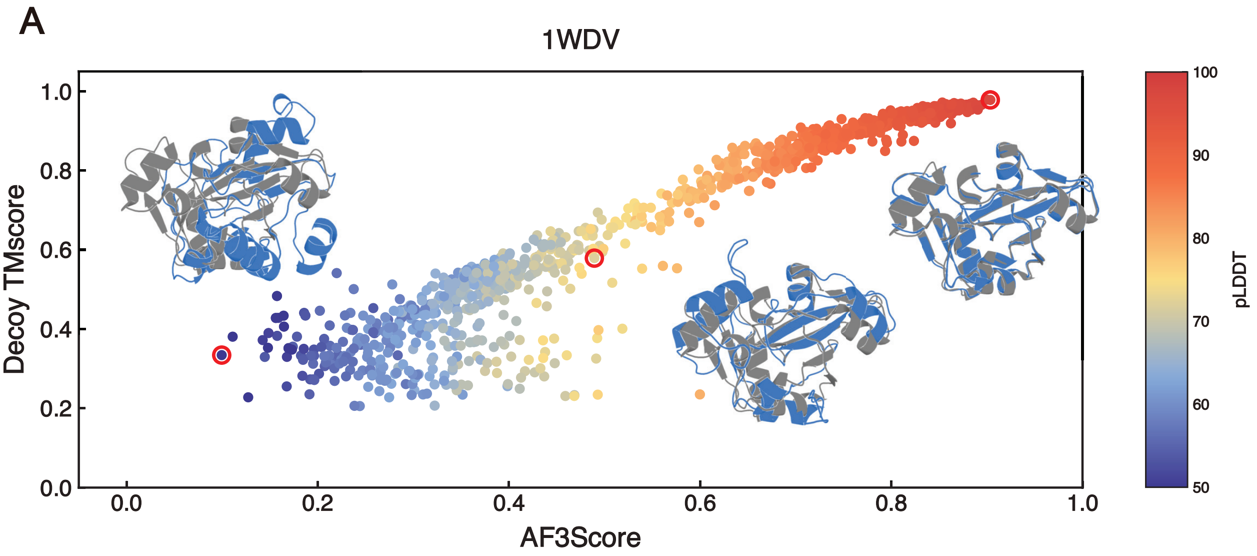


Figure S1: Demonstration of the correlation between scores generated by AF3Score and their corresponding TM-scores for target 1WDV. The structural superpositions showing native structure (grey) and decoys (blue). Points colored by pLDDT values.

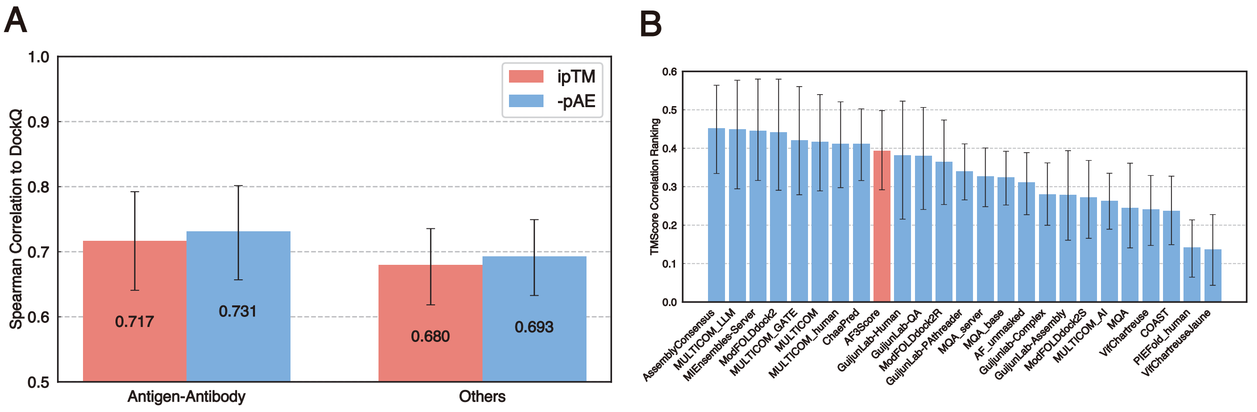


Figure S2: (A) Spearman correlations for antigen-antibody complexes (N=63) and other types of PPIs (N=125) in Docking Benchmark 5.5. Notably, the model demonstrated comparable performance on antigen-antibody complexes and non-AbAg PPIs (B) Rankings of assessment of global fold quality by 26 methods from CASP16.

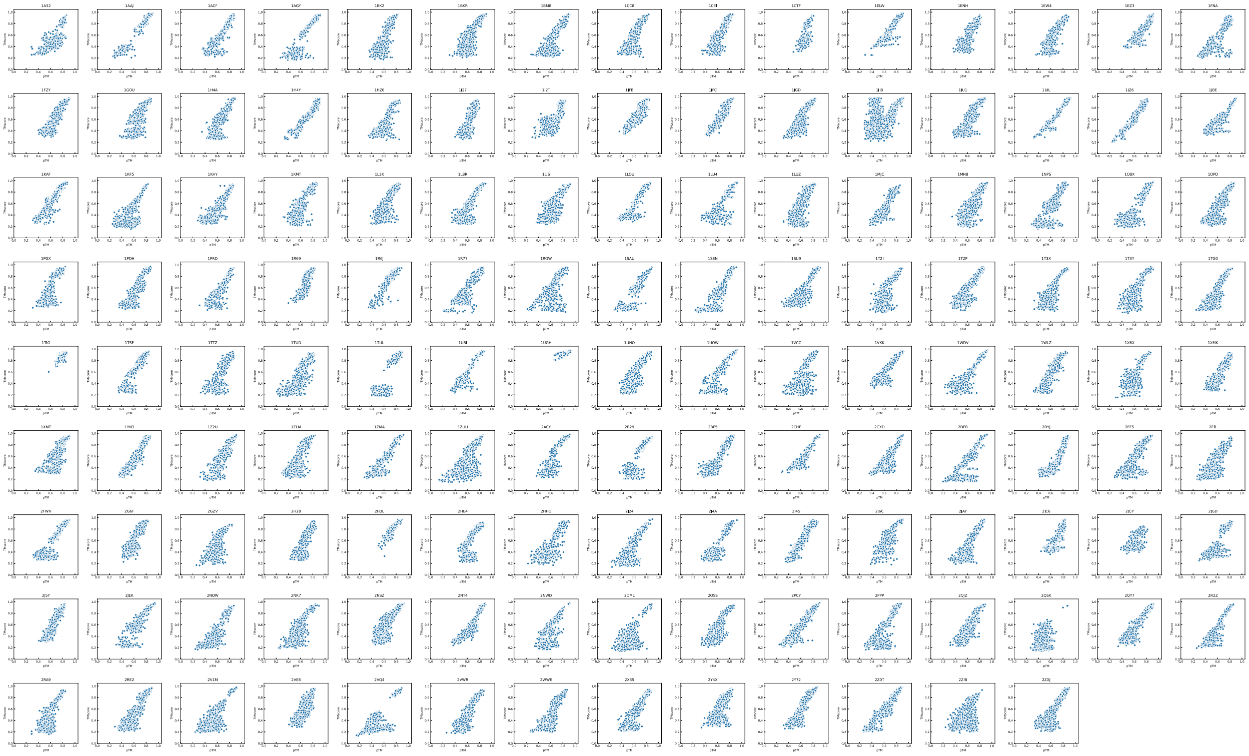

Figure S3: Correlation analysis between AF3Score predictions and TM-scores for all targets in the Rosetta Decoy dataset. Each subplot represents one target protein, showing the relationship between AF3Score values and decoy TM-scores. Points represent decoy structures.

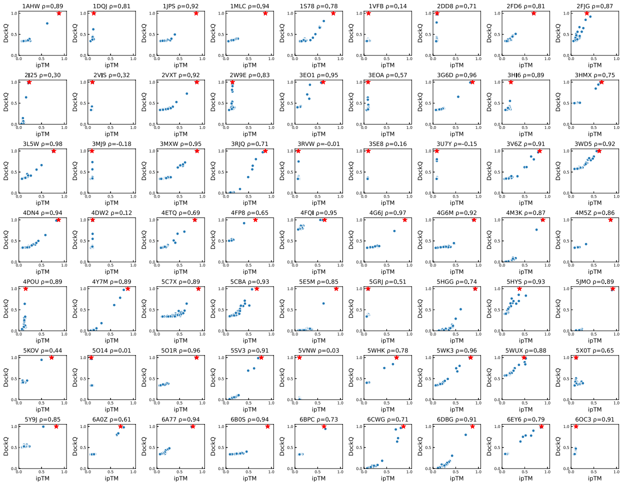


Figure S4: Correlation analysis between AF3Score ipTM and DockQ scores for antigen-antibody complexes from the Docking Benchmark 5.5 dataset. Each panel represents one complex with its PDB ID and Spearman correlation coefficient shown above. Native structures are marked with red stars.

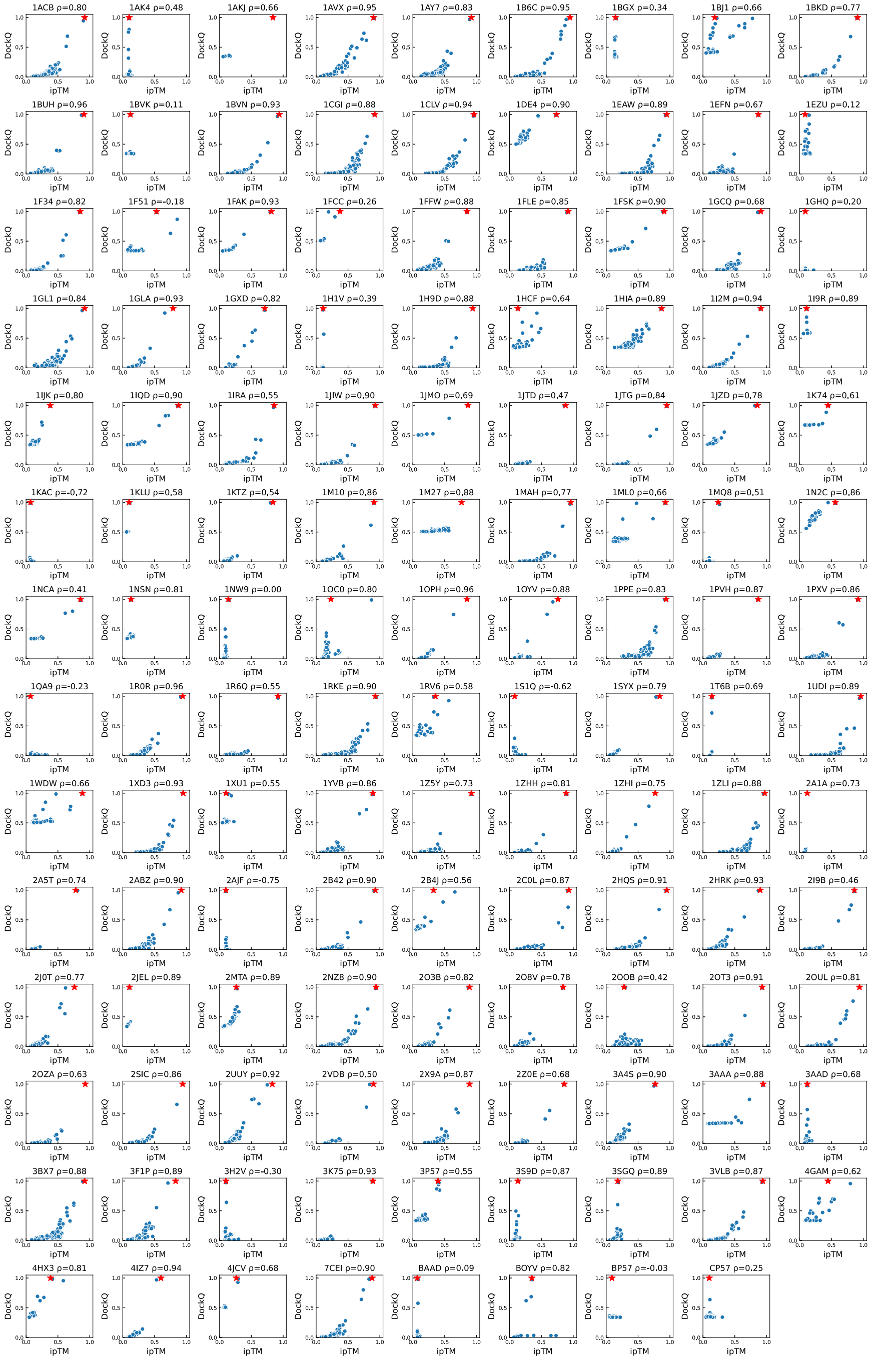

Figure S5: Correlation analysis between AF3Score ipTM and DockQ scores for non-antibody protein-protein complexes from the Docking Benchmark 5.5 dataset. Each panel represents one complex with its PDB ID and Spearman correlation coefficient shown above. Native structures are marked with red stars.

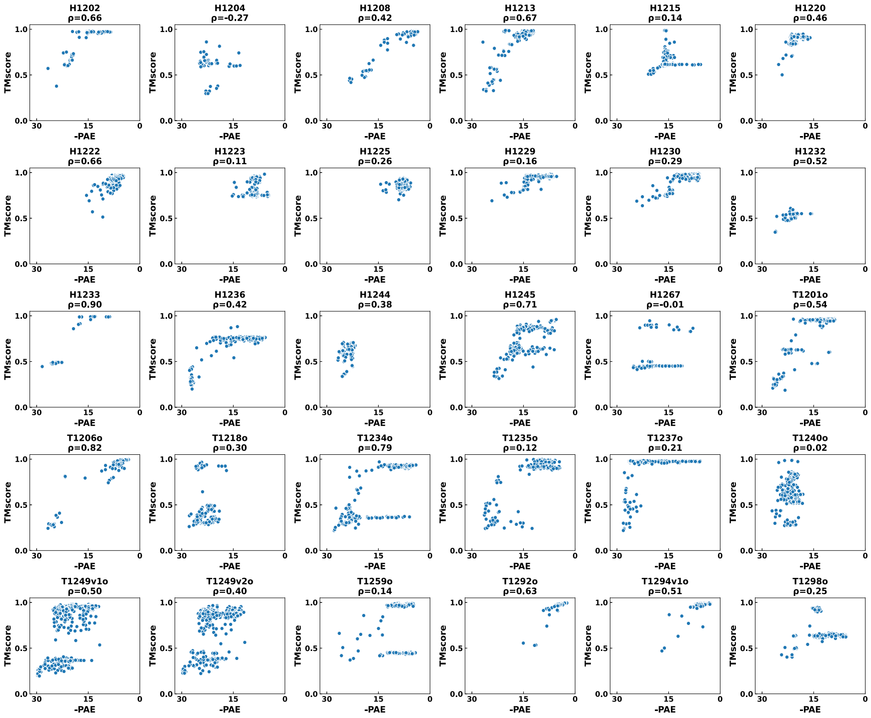


Figure S6: Correlation between predicted aligned error (pAE) and TMscore for CASP16 QMODE1 targets. Each subplot represents one target, with Spearman correlation coefficient shown in the title.

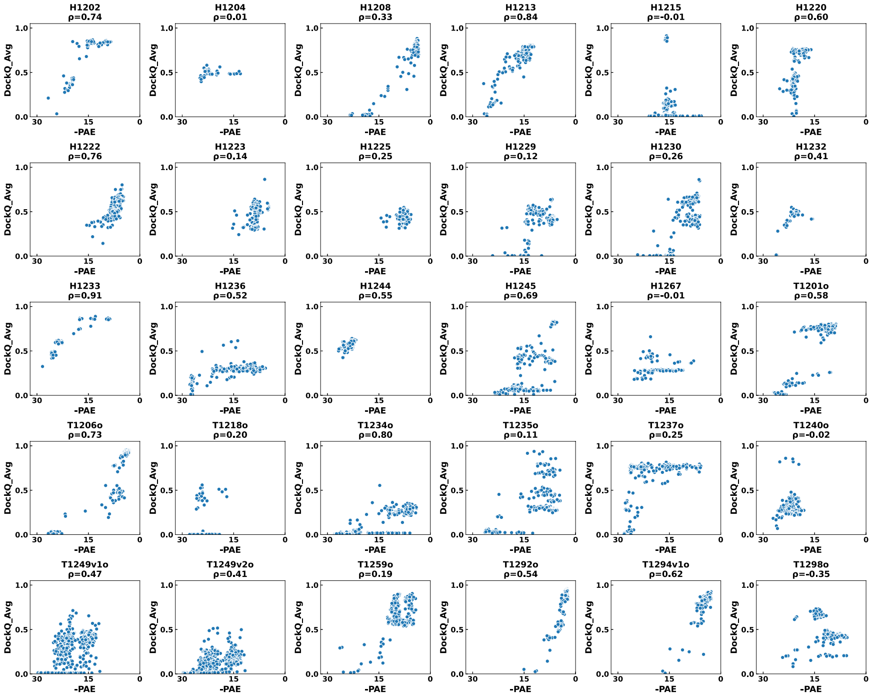


Figure S7: Correlation between predicted aligned error (pAE) and DockQ score for CASP16 QMODE1 targets. Each subplot shows the correlation for individual complexes, with Spearman correlation coefficient displayed in the title.

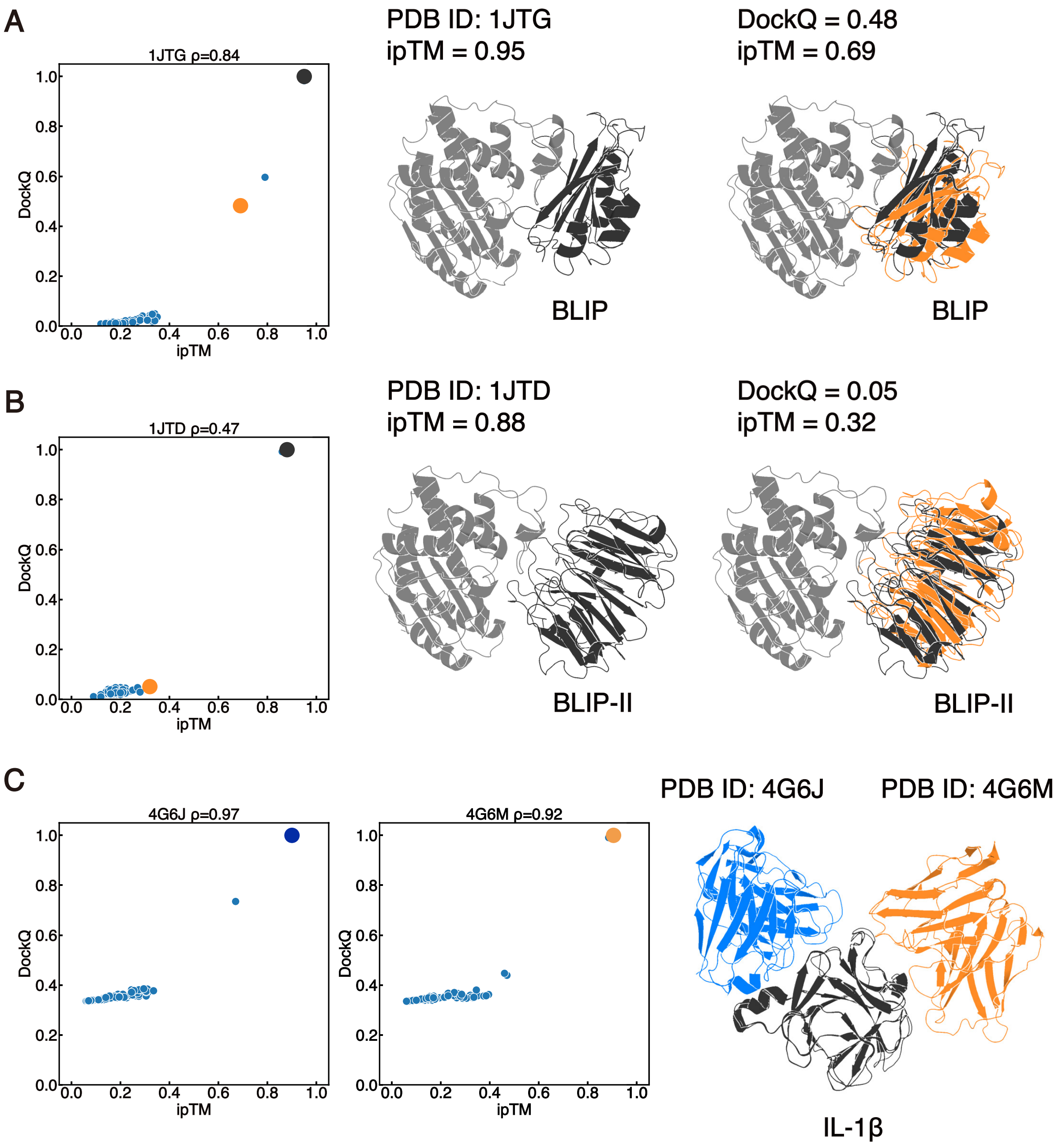


Figure S8: Case studies from the evaluations of AF3Score on PPIs. (A,B) Structural superposition of BLIP and BLIP-II complexes with TEM-1 $\beta$-lactamase (grey), showing native inhibitor structures (black) and decoy conformations (orange). Corresponding correlation plots display AF3Score values for native (black) and decoy (orange) structures. (C) Structural comparison of IL-1$\beta$ binding to two different antibodies with two distinct epitopes: canakinumab (PDB ID: 3G6J, blue) and gevokizumab (PDB ID: 3G6M, orange).

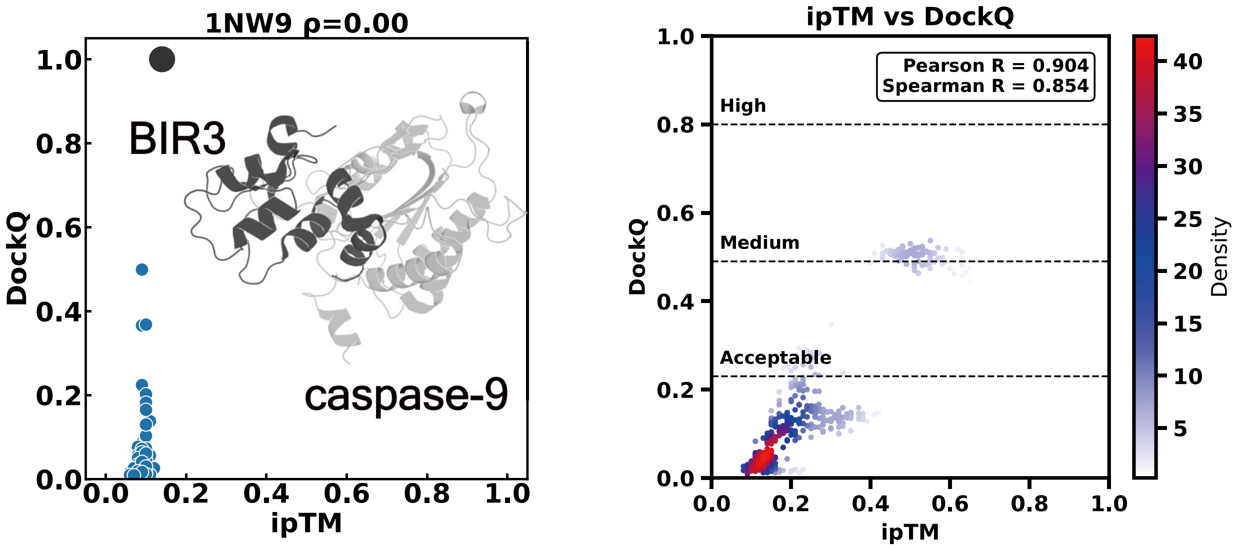


Figure S9: Analysis of a challenging case: the BIR3-caspase-9 complex. Left: Correlation between AF3Score ipTM and DockQ scores, with HDock-generated decoys shown in blue and the native structure in black. The structural representation shows the native complex with BIR3 in black and caspase-9 in grey. Right: Distribution of AF3-sampled structures (500 structures from 100 seeds, 5 structures per seed) plotted as ipTM versus DockQ scores, with sampling density indicated by color intensity. Horizontal lines mark quality thresholds for complex structures (Acceptable: 0.23, Medium: 0.49, High: 0.8).

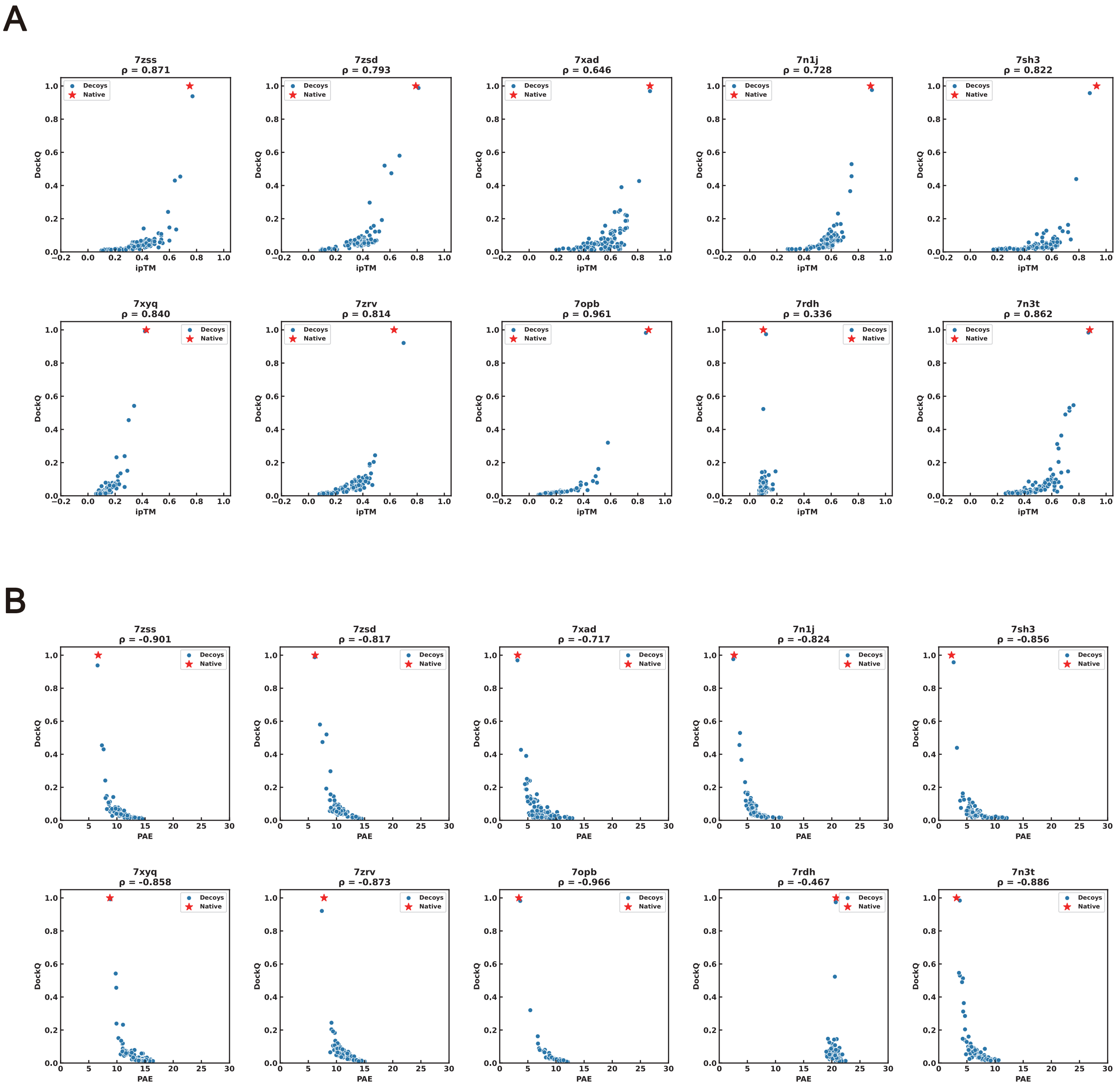


Figure S10: AF3Score evaluation of mini-binder decoys across 12 targets. To test the capability of identifying the correct binding interfaces among the decoy interfaces, structural decoys were generated from experimentally solved binder complexes. (a) Correlation between AF3Score ipTM and DockQ scores without MSA input. (b) Correlation between AF3Score ipTM and DockQ scores with AlphaFold2-searched MSA input. In both panels, red stars indicate native structures and blue dots represent HDOCK-generated decoys. Each subplot represents one target protein, with all structures using self-decoy as templates.

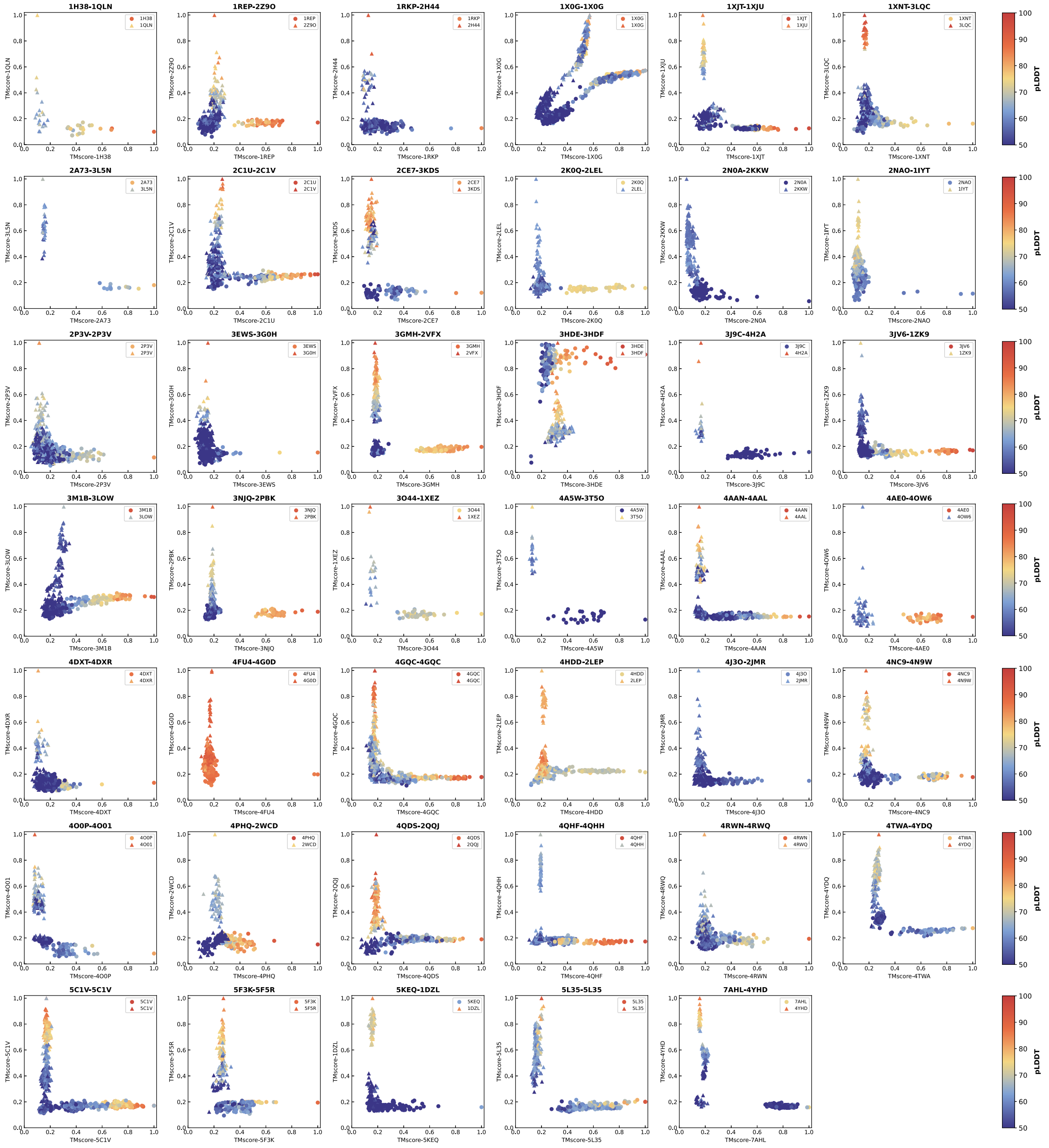


Figure S11: Comprehensive analysis of AF3Score evaluation on fold-switching protein decoys. Two-dimensional plots showing the relationship between TM-scores (relative to each native fold) and AF3Score confidence values for all analyzed fold-switching proteins. Each point represents a structure, with its color intensity indicating the CA pLDDT score of the fold-switching region as assessed by AF3Score. Different conformational states are distinguished by triangles and circles. The x and y axes represent TM-scores calculated against each of the two native folds, allowing visualization of structural similarity to both conformational states simultaneously. This analysis demonstrates AF3Score's ability to evaluate structures across the conformational landscape between the two native states.

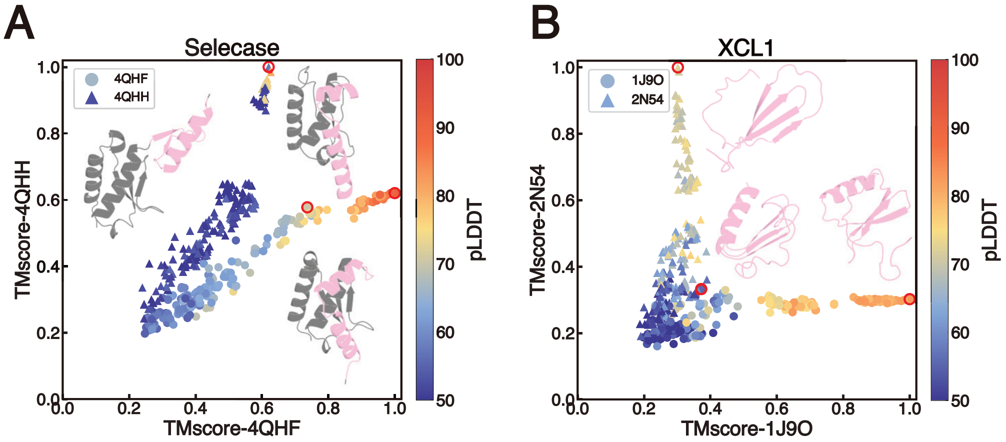


Figure S12: Analysis of the representative fold-switching proteins. The x- and y-axes represent TM-scores relative to each native fold. Each point corresponds to a structure, colored according to the C$\alpha$ pLDDT score of its fold-switching region. Triangles and circles distinguish between the two alternative folds. Native structures and selected decoys (highlighted by red circles) are shown, with fold-switching regions in pink and the remaining structure in grey.

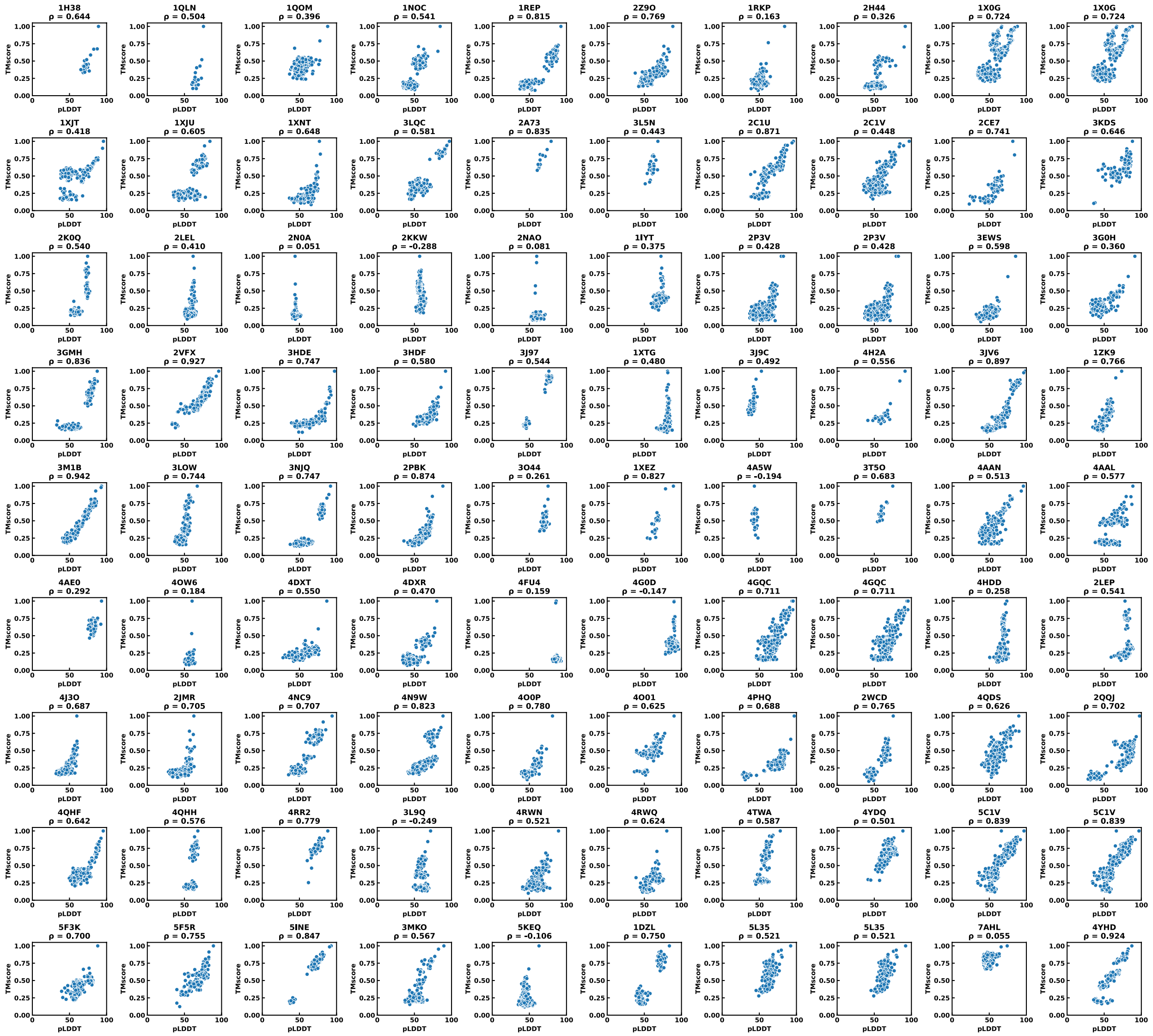


Figure S13: Correlation analysis between AF3Score pLDDT metrics and TM-scores for fold-switching proteins. Each subplot represents one fold-switching protein from our dataset, showing the relationship between AF3Score's predicted pLDDT values and TM-scores calculated against native conformations. Points represent individual decoy structures, with correlation coefficients shown for each case.

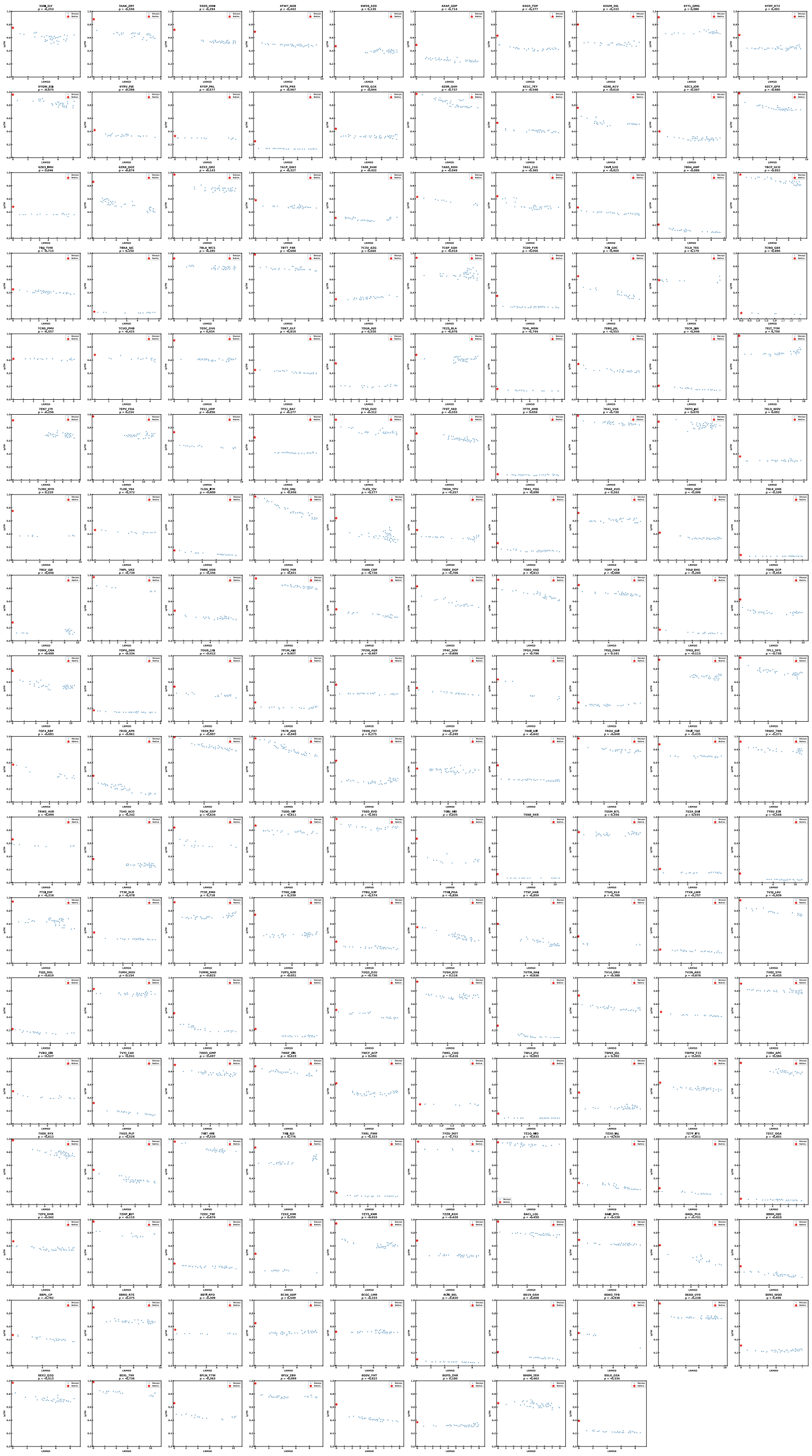

Figure S14: Correlation between ligand RMSD and AF3Score ipTM values for protein-ligand complexes from the PoseBusters dataset. Each subplot represents one complex, with docking-generated decoys shown in blue and native structures in red. The x-axis shows ligand RMSD values, and the y-axis shows AF3Score ipTM scores.

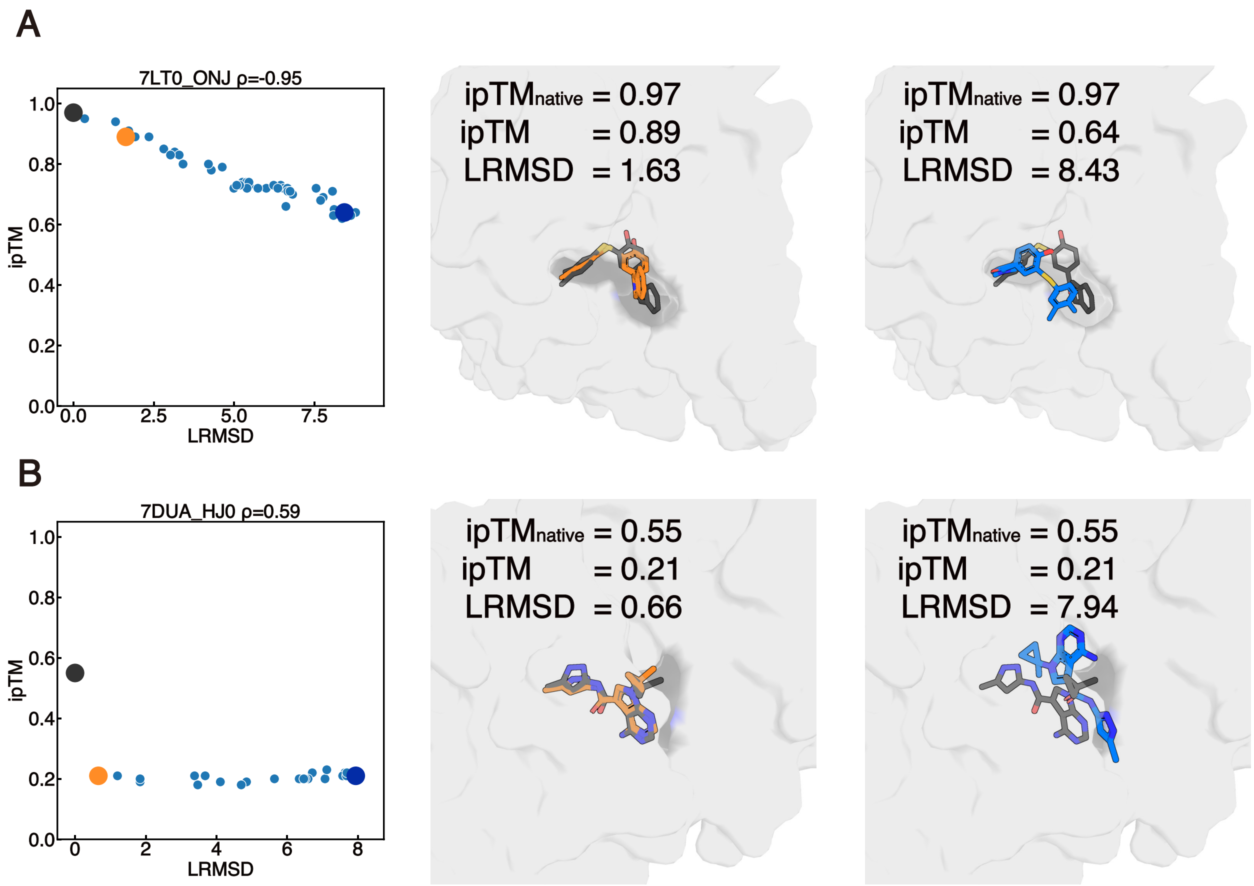


Figure S15: Case studies of the evaluation of AF3Score on the protein-ligand complex dataset. (A,B) Views of ligand interactions with protein pockets (grey surface) for 7LT0 and 7DUA complexes. Correlation plots show AF3Score's ipTM versus LRMSD values, with representative structures highlighting near-native (yellow) and high-LRMSD (blue) poses. While AF3Score correctly identified the native pose (ipTM = 0.55), it failed to properly rank near-native decoys. Specifically, poses with LRMSD of 0.66Å received the same low scores (ipTM = 0.21) as clearly incorrect poses (LRMSD = 7.94Å)
